# Human induced pluripotent stem cell-derived microglia contribute to the pathophysiology of Fragile X syndrome via increased RAC1 signaling

**DOI:** 10.1101/2024.06.24.600387

**Authors:** Poulomi Banerjee, Shreya Das Sharma, Karen Burr, Kimberley Morris, Tuula Ritakari, Paul Baxter, James D Cooper, Alessandra Cardinalli, Srividya Subash, Evdokia Paza, David Story, Sumantra Chattarji, Peter C Kind, Neil O Carragher, Bhuvaneish T Selvaraj, Josef Priller, Siddharthan Chandran

## Abstract

Fragile X syndrome (FXS) is one of the most common monogenic causes of neurodevelopmental disorders characterized by intellectual disability, autism and epilepsy. Emerging evidence suggests a role for immune dysfunction in autism. Using induced pluripotent stem cell (iPSC)-derived microglial cells from FXS patients (mFXS-MG) and *FMR1*-deficient microglia from *FMR1*-knock out human embryonic stem cells (*FMR1 KO*-MG), we show that loss-of-function of Fragile X Messenger Ribonucleoprotein (FMRP) leads to cell autonomous phagocytic deficits and a proinflammatory state in microglia when compared to gene-corrected controls. Moreover, increased RAC1 signaling in mFXS-MG and *FMR1 KO*-MG results in increased actin polymerization and enhanced activation of NF-κB signaling. Exposure of control iPSC-derived cortical neuron cultures to conditioned medium from proinflammatory mFXS-MG results in hyperexcitability. Importantly, pharmacological inhibition of RAC1 signaling in mFXS-MG attenuates their proinflammatory profile and corrects the neuronal hyperexcitability caused by the conditioned medium. Our results suggest that microglia impair neuronal function in FXS, which can be prevented by targeting of RAC1 signaling.

**Significance statement:** FXS is one of the most common monogenic causes of neurodevelopmental disorders characterized by intellectual disability, autism, epilepsy and has been associated with immune dysfunction. We therefore generated brain macrophages (microglia) from patient-derived induced pluripotent stem cells (mFXS-MG) and an embryonic stem cell line deficient in the Fragile X messenger ribonucleoprotein 1 (*FMR1* KO-MG). We find enhanced activation of RAC1 signaling resulting in phagocytic deficits and immune activation of mFXS-MG and *FMR1* KO-MG. Exposure of control iPSC-derived cortical neurons to conditioned medium from proinflammatory mFXS-MG results in neuronal hyperexcitability, which can be prevented by pharmacological RAC1 inhibition in mFXS-MG. We conclude that RAC1 signaling in microglia could be a potential therapeutic target in FXS.

## Introduction

FXS is one of the most common monogenic causes of neurodevelopmental disorders characterized by intellectual disability, autism and epilepsy. FXS is caused by a CGG repeat expansion in the promoter region of the Fragile X messenger ribonucleoprotein 1 (*FMR1*) gene, resulting in the epigenetic silencing of the *FMR1* gene and subsequent loss of its protein product, Fragile X Messenger Ribonucleoprotein (FMRP) (1, 2).

Whilst the pathophysiology of autism remains poorly understood, emerging evidence suggests a potential role for dysregulated circulating myeloid cells and brain resident macrophages/microglia contributing to disease (3). Bulk transcriptome data, single cell sequencing data and positron emission tomography (PET) studies show microglial activation (4–7) in brain tissue samples of people with autism. Studies of blood samples from people with autism have also reported elevated levels of proinflammatory cytokines. Together these findings highlight immune dysfunction in patients with autism (8, 9). Treatment of preclinical animal models with bacterial lipopolysaccharide or cytokines, such as IL-6 also provide indirect evidence for microglial cells contributing to autistic behaviours (10, 11). Interestingly, selective depletion of Transmembrane protein 59 (TMEM59) from microglia in mice, a gene known to be associated with autism, resulted in cell autonomous phagocytic deficits in murine microglia and non-cell autonomous neuronal dysfunction such as enhanced excitatory synaptic transmission and autism-like behavioural deficits (12). Moreover, Drosophila melanogaster *Fmr1* mutants also exhibit increased vulnerability to bacterial infection and reduced phagocytosis of bacteria by systemic immune cells (13). Importantly, immune dysregulation has been observed in blood samples of FXS patients and systematic comorbidity analyses have revealed increased incidence of infectious diseases in FXS patients (14, 15).

A key molecular regulator of innate immune cell and microglial functions, including phagocytosis, is Ras-related C3 botulinum toxin substrate 1 (RAC1) signaling. Specifically, activated RAC1 signaling regulates the dynamics of the actin cytoskeleton through proteins such as COFILIN during phagocytosis (16). COFILIN is an actin depolymerizing factor which can sever filamentous actin (F actin) and generate new actin monomers (G actin) (17, 18). However, sustained activation of RAC1 signaling leads to inactivation of COFILIN, which results in an increase of F actin and causes reduced phagocytosis (19). Separately, RAC signaling can regulate nuclear factor-κB (NF-κB), which is an ubiquitously expressed transcription factor that plays a central role in regulating microglial immune activation (20, 21). Activated RAC1 signaling can positively regulate NF-κB signaling through downstream PI3K, AKT pathway and the expression of dominant-negative RAC1 can block NF-κB transactivation (22).

Against this background, it is of considerable interest that FMRP interacts directly with Cytoplasmic FMRP interacting protein 1(CYFIP1) which is a key binding partner of RAC1-GTP (23). Specifically, CYFIP1 shuttles across two protein complexes-1) a translation-repressive machinery comprised of FMRP and 2) a cytoskeletal RAC1-WAVE regulatory complex. In the absence of FMRP, the equilibrium of these complexes changes and interactions of CYFIP1 and RAC1-WAVE are favoured which leads to enhanced activation of actin cytoskeleton remodelling pathway (24). Given that, microglial cells are immune cells which undergo changes in actin polymerisation during phagocytosis, we decided to explore the role of RAC1 signaling in microglial cells lacking FMRP and the potential implications for the pathophysiology of FXS.

## Results

### *Loss-of-function* of FMRP leads to impaired phagocytosis in mFXS-MG and *FMR1*-KO MG through actin polymerisation deficits

Isolated monocultures of iPSC-derived microglial cells were generated from two FXS patients (mFXS-1, mFXS-2), gene-edited isogenic controls (isoFXS-1, isoFXS-2), and control iPSCs (CTRL-iPSC) employing an established protocol (25, 26) (Suppl Figs 1, 2A-D). Likewise, microglial cells were generated from *FMR1*-/- human embryonic stem cells (*FMR1* KO) and control human embryonic stem cells (CTRL-ES). Microglial cells with diseased genotypes (mFXS-1, mFXS-2, *FMR1* KO MG) and control genotypes (isoFXS-1, isoFXS-2, CTRL (iPSC), CTRL(ES) MG) exhibited no significant difference in the expression of microglial key marker genes such as *TMEM119* and *SALL1* (Suppl. Figs 2E). As expected, loss of FMRP was observed in mFXS MG, *FMR1*-KO MG compared to isoFXS MG and CTRL (ES) MG (Fig 1A). To assess the impact of loss-of-function of FMRP on microglial phagocytosis, we quantified time-dependent internalisation of pH-conjugated zymosan bioparticles, which revealed significantly fewer internalised bioparticles in mFXS MG and *FMR1*-KO MG compared to isoFXS MG and CTRL(ES) MG suggesting a phagocytic deficit (Fig 1B, Supplementary movies 1-3).

**Figure 1:**
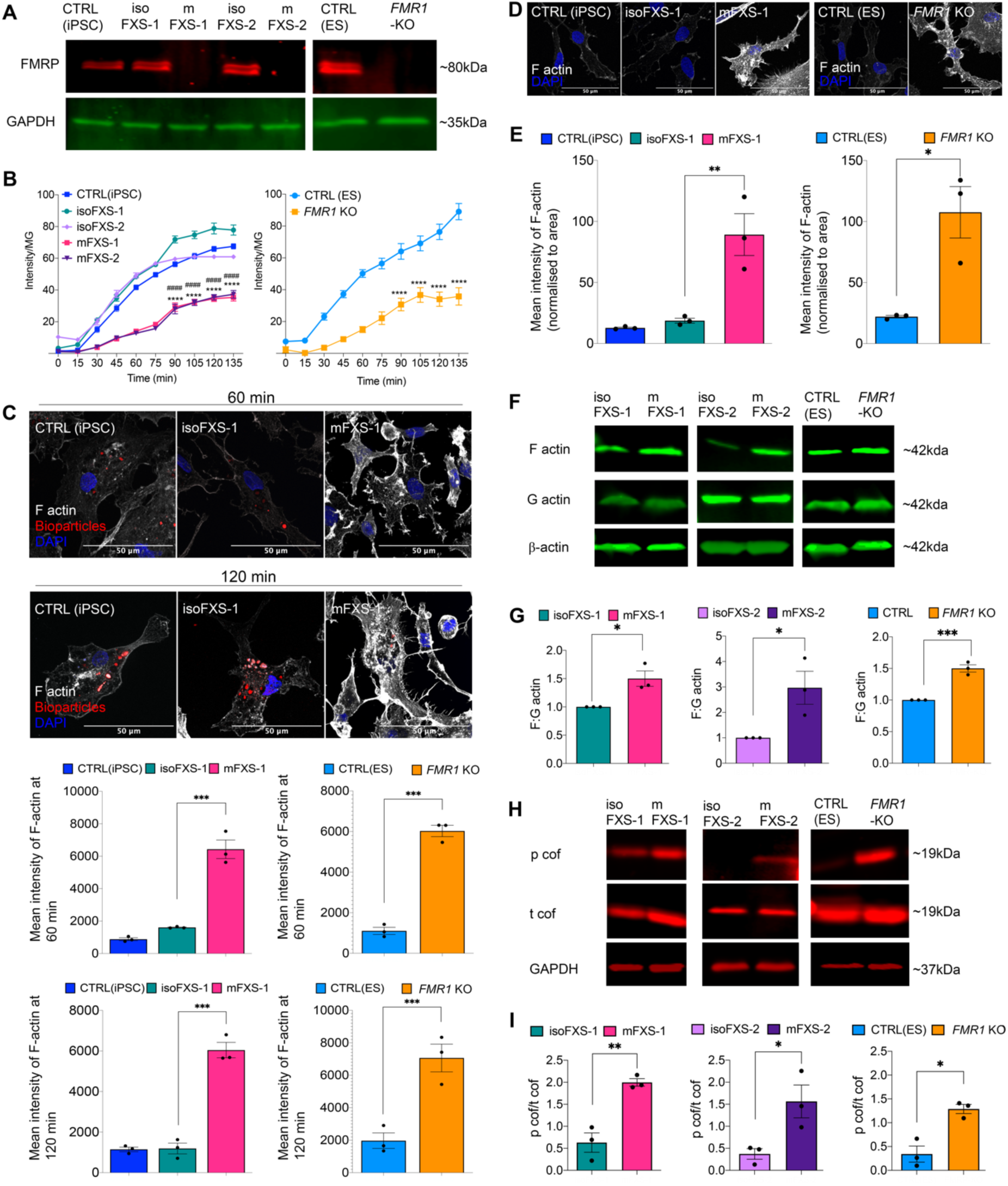
Aberrant actin polymerization underlies phagocytic deficit in mFXS-MG and FMR*1* KO-MG. **(A)** Western blot depicting loss of FMRP in mFXS-MG and *FMR1* KO-MG **(B)** Graphs showing real-time imaging of zymosan bioparticle uptake at 15 min intervals, demonstrating a phagocytic deficit in mFXS 1-MG, mFXS 2-MG (left) and *FMR1* KO-MG (right). Statistical analysis was performed across isoFXS 1-MG, mFXS 1-MG, isoFXS 2-MG, mFXS 2-MG and CTRL(ES)-MG, *FMR1* KO-MG using two-way ANOVA and Tukey’s multiple comparison test, data are represented as mean +/- SEM; N=3 (**** p < 0.0001 represents the value for mFXS-1 and isoFXS-1 and ####p<0.0001 represents the value for mFXS-2 and isoFXS-2). **(C)** Representative images of immunofluorescence staining with phalloidin showing aberrant F actin during phagocytosis of zymosan bioparticles (red) in mFXS 1-MG at 60-minutes (top panel) and 120-minutes (bottom panel). Graphs showing increased mean fluorescent intensity (pixel intensity normalised to cell area) of F actin during phagocytosis at 60-minute (top row) and 120-minute (bottom row) timepoint for mFXS 1-MG and *FMR1* KO-MG when compared to isoFXS 1-MG and CTRL (ES)-MG. Each dot represents averaged data from one biological replicate comprising of 6-8 cells, Data represent mean ± SEM from three biological replicates (N=3), statistical analysis was performed using one way ANOVA with post-hoc Bonferroni’s test (***p<0.001) **(D)** Representative images of immunofluorescence staining with phalloidin showing aberrant F actin in mFXS 1-MG and *FMR1* KO-MG at a basal state **(E)** Graphs represent increased mean fluorescent intensity (pixel intensity normalised to cell area) of F actin at basal state in mFXS 1-MG (left) and *FMR1* KO-MG (right) when compared to isoFXS1-MG and CTRL(ES)-MG. Each dot represents averaged data from one biological replicate comprising of 6-8 cells. Data represent mean ± SEM from three biological replicates (N=3), statistical analysis was performed using one way ANOVA with post-hoc Bonferroni’s test (* p < 0.05,** p <0.01). **(F)** Western blot depicting F-actin and G actin in isoFXS 1-MG, mFXS 1-MG, isoFXS 2-MG, mFXS 2-MG, CTRL(ES)-MG and *FMR1* KO-MG **(G)** Graph showing relative increase of F/G actin ratio in mFXS 1-MG, mFXS 2-MG and *FMR1* KO-MG when compared to isoFXS 1-MG, isoFXS 2-MG and CTRL(ES)-MG as determined by densitometric analysis of immunoblot scans after normalization with beta-actin (loading control). Each dot represents data from one biological replicate. Data represent mean ± SEM from three independent experiments N=3, statistical significance was calculated across isoFXS 1-MG, mFXS 1-MG and isoFXS 2-MG, mFXS 2-MG and CTRL(ES)-MG, *FMR1* KO-MG using unpaired T test (* p < 0.05 \*\*\**P*<0.001) **(H)** Western blot for total COFILIN and phospo-COFILIN in isoFXS 1-MG, mFXS 1-MG, isoFXS 2-MG, mFXS 2-MG, CTRL(ES)-MG and *FMR1* KO-MG **(I)** Graph showing relative increase of phospo-COFILIN to total COFILIN ratio in mFXS 1-MG, mFXS 2-MG, *FMR1* KO-MG when compared to isoFXS 1-MG, isoFXS 2-MG and CTRL(ES)-MG as determined by densitometric analysis of immunoblot scans. Each dot represents data from one biological replicate. Data represent mean ± SEM from three independent experiments N=3, statistical significance was calculated across isoFXS 1-MG, mFXS 1-MG and isoFXS 2-MG, mFXS 2-MG and CTRL(ES)-MG, *FMR1* KO-MG using unpaired T Test (* p < 0.05, **p<0.01) N represents number of times experiments were performed using cells generated from independent iPSC differentiations.

Given that, the dynamics of actin polymerisation and depolymerisation are crucial during phagocytosis and absence of FMRP can prompt increased actin polymerisation (19, 24), we investigated the level of polymerised actin (F actin) following a challenge to internalise zymosan bioparticles (Fig 1C). Phalloidin staining revealed that both mFXS MG and *FMR1* KO MG have significantly increased F actin at 60 and 120-minute time point when compared to isoFXS MG and CTRL (ES) MG (Fig 1C, Suppl. Fig 3). Indeed mFXS-MG and *FMR1* KO MG showed significantly increased F actin when they were not challenged with zymosan bioparticles, suggestive of a cell autonomous deficit in actin dynamics in microglial cells lacking FMRP (Figs 1D, 1E). To further quantify and validate the dynamics of actin polymerisation and depolymerisation, we determined the ratio of filamentous (F) *vs*. globular (G) actin through protein fractionation in the control and FXS microglial cells. These experiments revealed that mFXS MG and *FMR1* KO MG have significantly increased F-actin:G-actin ratios compared to isogenic MG and CTRL(ES) MG (Figs. 1F, 1G). Given that, COFILIN is an actin-depolymerizing factor and that inactivation of COFILIN by phosphorylation could lead to an increase in actin polymerisation (18), we next examined the abundance of COFILIN and its phosphorylation status in FXS and control microglial cells. Phosphorylation of COFILIN was significantly increased in both mFXS MG and *FMR1*-KO MG microglial cells, while total COFILIN was not altered across control and disease genotypes (Fig 1H, 1I), indicating that inactivation of COFILIN is associated with increased actin polymerization (Fig 1C-1G, Suppl. Fig 3) in mFXS-MG and *FMR1* KO-MG. Collectively, these findings suggest that the absence of FMRP in microglial cells leads to impaired phagocytosis of zymosan bioparticles due to altered actin dynamics.

### *Loss-of-function* of FMRP in microglial cells leads to a proinflammatory state in mFXS MG and *FMR1* KO MG through activation of the NF-κB pathway

We next assessed the activation state of mFXS-MG and *FMR1* KO-MG by performing qPCR for the microglial homeostatic marker gene *P2RY12* and a proinflammatory cytokine gene *1L-1b* and also quantifying levels of a panel of cytokines released in the secretomes of FXS versus control microglial cells at the basal level by forward phase cytokine assay. qPCR data revealed significant downregulation of microglial homeostatic gene *P2Y12* mRNA expression and a corresponding significant upregulation of *IL1b* mRNA expression in mFXS MG and *FMR1* KO MG (Figs. 2A, 2B). qPCR data was further corroborated with a cytokine array, which revealed increased levels of the proinflammatory cytokines RANTES, IL-1a, IL-1b, TNF-a in both mFXS MG and *FMR1* KO MG, indicating a cell intrinsic proinflammatory nature of microglial cells lacking FMRP (Figs. 2C, 2D). Given that the NF-κB pathway is the master regulator for immune activation in microglia (21), we next assessed the activation status of NF-κB in FXS and control microglial cells. Both mFXS MG and *FMR1* KO MG displayed a significantly increased localisation of NF-κB in the nucleus compared to isoFXS MG and CTRL(ES) MG, indicating increased activation of the NF-κB pathway in microglial cells lacking FMRP (Figs. 2E-2H). Collectively, these data suggest that the absence of FMRP in microglial cells leads to activation of the NF-κB pathway which results in an increased production of proinflammatory cytokines.

**Fig 2:**
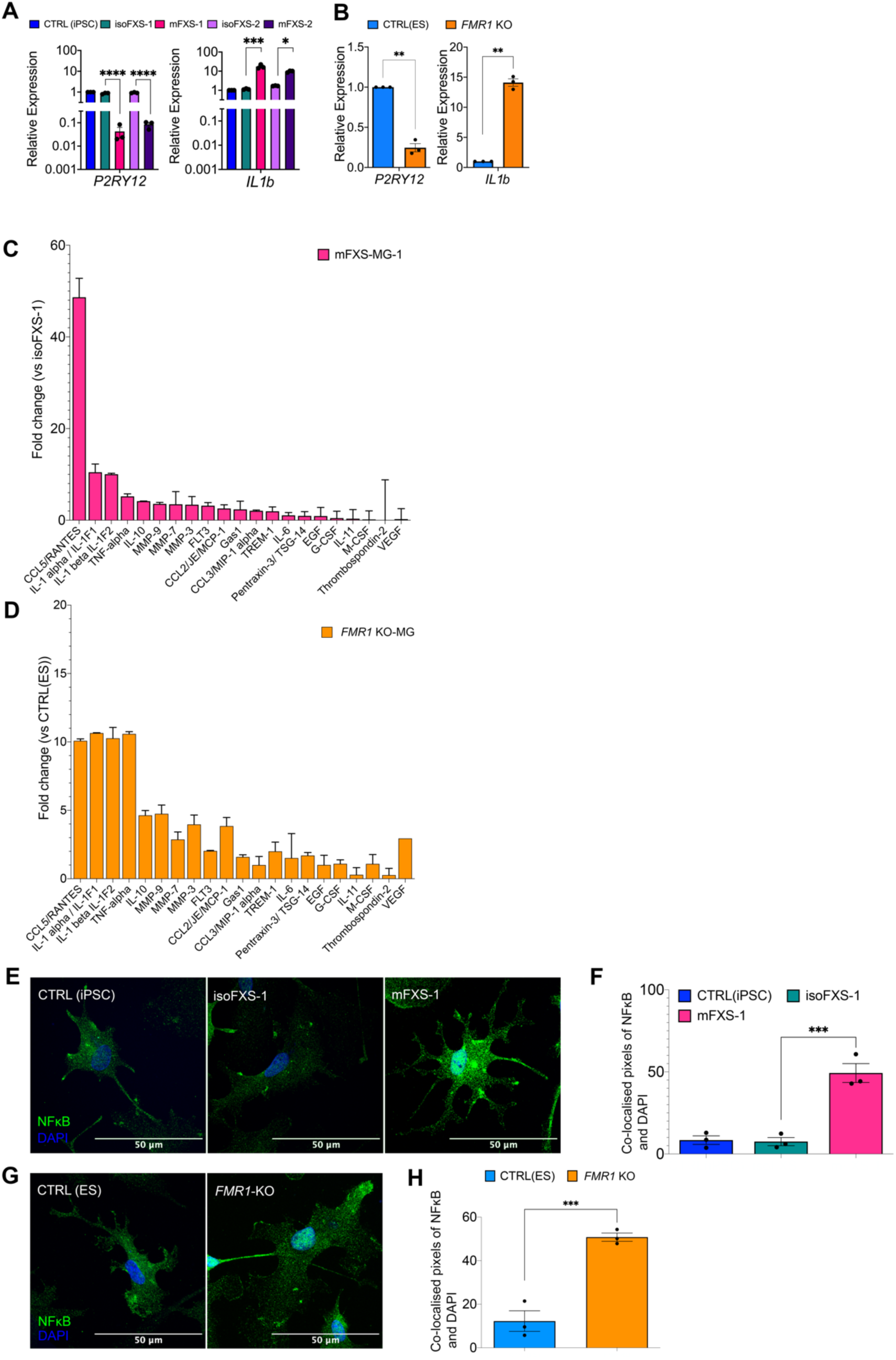
Activated NFκB pathway drives a pro-inflammatory profile in mFXS-MG and *FMR1* KO-MG. **(A,B)** Bar graphs representing reduced expression of *P2RY12* and increased expression of *IL1b* in mFXS 1-MG, mFXS 2-MG **(A)**, and *FMR1* KO-MG **(B)**. Each dot represents data from one biological replicate. Data represent mean ± SEM from three biological replicates (N=3). Statistical analysis was performed across isoFXS 1-MG, mFXS 1-MG, isoFXS 2-MG, mFXS 2-MG using one-way ANOVA and Bonferroni’s multiple comparison test (* p < 0.05, *** p <0.001, **** p <0.0001) and CTRL(ES)-MG, *FMR1* KO-MG using paired T test (** p < 0.01) **(C,D)** Bar graphs demonstrating top 21 commonly dysregulated cytokines in mFXS 1-MG **(C)** and *FMR1* KO-MG **(D).** Y axis represents the fold change versus their isogenic controls. **(E,G)** Representative immunofluorescence staining showing enhanced nuclear localisation of NFκB suggestive of active NFκB signaling in mFXS 1-MG and *FMR1* KO-MG when compared to isoFXS 1-MG and CTRL (ES)-MG **(F,H)** Graphs showing increased colocalised pixels of NFκB and DAPI in mFXS 1-MG and *FMR1* KO-MG when compared to isoFXS 1-MG and CTRL (ES)-MG. Each dot represents averaged data from one biological replicate comprising 6-8 cells. Data represent mean ± SEM from three biological replicates (N=3). Statistical analysis was performed across isoFXS 1-MG, mFXS 1-MG, and CTRL(ES)-MG, *FMR1* KO-MG using one way ANOVA with post-hoc Bonferroni’s test (***p<0.001) N represents number of times experiments were performed using cells generated from independent iPSC differentiations.

### Activation of RAC1 signaling in mFXS-MG and *FMR1*-KO MG mediates aberrant actin polymerisation and activation of NF-κB signaling

As the RAC1 signaling pathway can regulate both actin cytoskeletal dynamics (16) and NF-κB signaling (20), we next examined RAC1 activity in diseased and control microglial cells through PAK-PBD pull down. The RAC/CDC42 (p21) binding domain (PBD) of the human p21 activated kinase 1 (PAK) protein binds specifically to GTP-bound RAC1 protein (activated RAC1), therefore GST-PAK fusion protein was used to specifically precipitate active GTP-bound RAC1. We found increased levels of activated RAC1 in mFXS MG and *FMR1*-KO MG when compared to isoFXS MG and CTRL (ES) MG with no change in the level of total RAC1 (Figs 3A, 3B). These findings indicate increased activation of RAC1 signaling in both mFXS MG and *FMR1*-KO MG.

**Fig 3:**
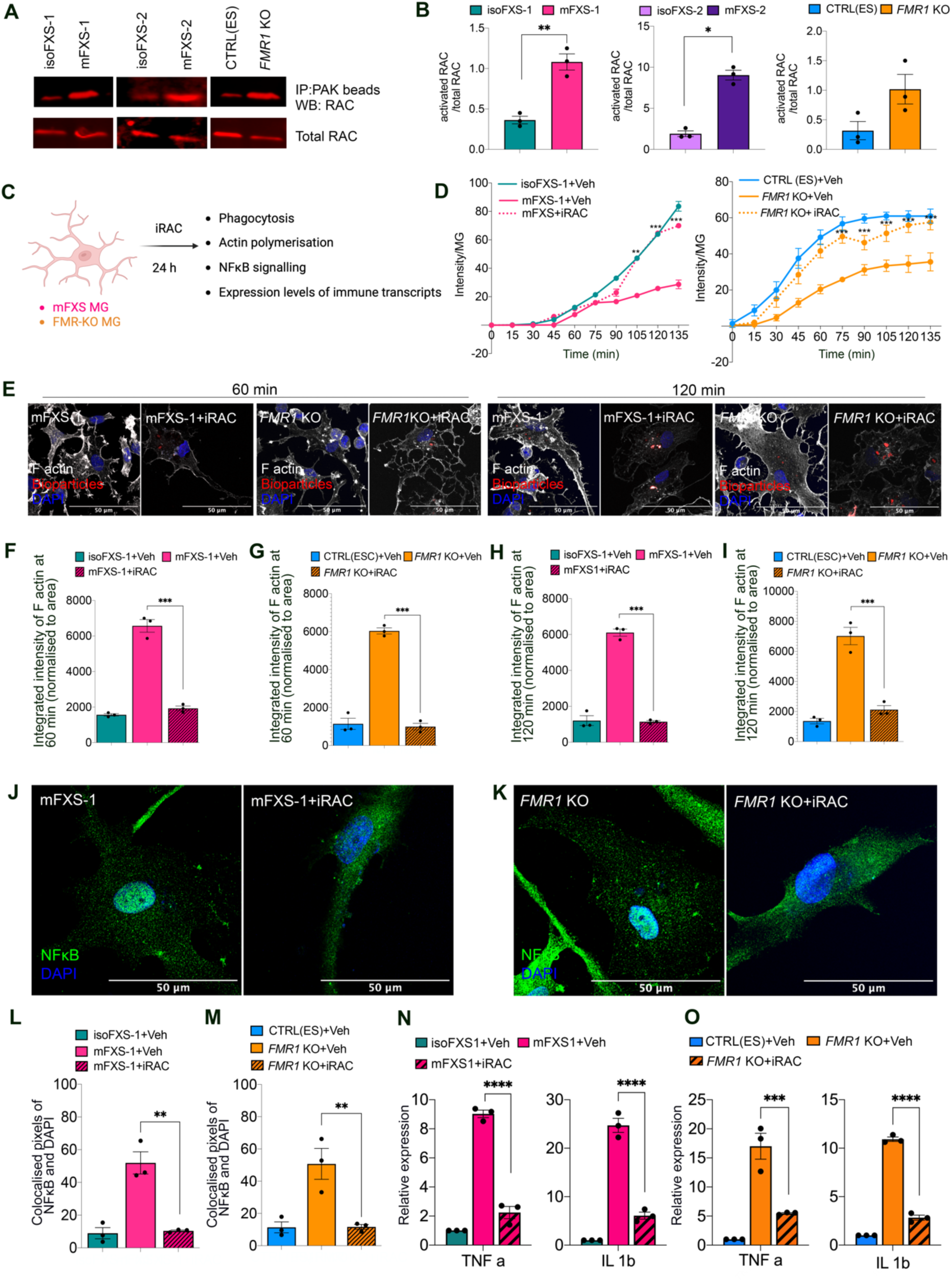
Loss of FMRP drives enhanced activation of RAC signaling and pharmacological inhibition of hyperactive RAC signaling ameliorates phagocytic defect and immune dysfunction in mFXS-MG and *FMR1* KO-MG. **(A)** Western blots for activated RAC pull down with PAK-PBD beads in mFXS1-MG, isoFXS1-MG, mFXS2-MG, isoFXS2-MG and *FMR1* KO-MG, CTRL(ES)-MG **(B)** Graphs for densitometric quantification of activated RAC and total RAC show enhanced activation of RAC signaling in mFXS1-MG, mFXS 2-MG and *FMR1* KO-MG. Each dot represents data from one biological replicate. Data represent mean ± SEM from three biological replicates (N=3), statistical significance was calculated across isoFXS 1-MG, mFXS 1-MG and isoFXS 2-MG, mFXS 2-MG and CTRL(ES)-MG, *FMR1* KO-MG using unpaired T Test (* p < 0.05, ** p < 0.01) **(C)** Graphic showing experimental paradigm for treating mFXS 1-MG and *FMR1* KO-MG with RAC inhibitor (iRAC) for 24 hours to assess phagocytosis, actin polymerisation, NFκB signaling pathway and expression of immune transcripts. **(D)** Graphs showing real-time imaging of zymosan bioparticle uptake at 15 min intervals, demonstrating a reversal of phagocytic deficit in mFXS1-MG, *FMR1* KO-MG following iRAC treatment (see dotted lines). Statistical analysis was performed across iRAC treated mFXS1-MG and untreated mFXS1-MG, and iRAC treated *FMR1* KO-MG and untreated *FMR1* KO-MG using two-way ANOVA and Bonferroni’s multiple comparison test, data are represented as mean +/- SEM; N=3 (***p<0.001). **(E)** Representative immunofluorescence images showing reduced F actin during phagocytosis at 60 minutes and 120 minutes in mFXS1-MG and *FMR1* KO-MG following treatment with iRAC. **(F-I)** Graphs showing reduced mean fluorescent intensity (pixel intensity normalised to cell area) of F actin during phagocytosis at 60-minute and 120-minute timepoint in iRAC treated mFXS1-MG and *FMR1* KO-MG when compared to untreated mFXS1-MG and *FMR1* KO-MG. Each dot represents averaged data from one biological replicate comprising 6-8 cells. Data represent mean ± SEM from three independent biological replicates (N=3), statistical analysis was performed using one way ANOVA with post-hoc Bonferroni’s test (***p<0.001) **(J,K)** Representative immunofluorescence staining showing reduced nuclear localisation of NFκB suggestive of reduced activation of NFκB signaling in iRAC treated mFXS 1-MG **(J)** and *FMR1* KO-MG **(K)** compared to untreated mFXS 1-MG and *FMR1* KO-MG. **(L,M)** Graphs showing reduced colocalised pixels of NFκB and DAPI in iRAC treated mFXS 1-MG **(L)** and *FMR1* KO-MG **(M)** compared to untreated mFXS 1-MG and *FMR1* KO-MG. Each dot represents data from one biological replicate comprising of 6-8 cells. Data represent mean ± SEM from three biological replicates (N=3), statistical analysis was performed using one-way ANOVA with post-hoc Bonferroni’s test (** p <0.01) **(O)** Graphs representing reduced expression of *TNFa* and *IL-1b*in iRAC treated mFXS1-MG **(left)** and *FMR1* KO-MG **(right)** compared to untreated mFXS1-MG and *FMR1* KO-MG. Each dot represents data from one biological replicate. Data represent mean ± SEM from three biological replicates (N=3), statistical analysis was performed using one way ANOVA with post hoc Tukey’s multiple comparison test (***p<0.001, **** p <0.0001). N represents number of times experiments were performed using cells generated from independent iPSC differentiations.

Based on literature (16, 20), we hypothesized that activated RAC1 signaling could lead to increased actin polymerisation and enhanced activation of the NF-κB pathway in mFXS MG and *FMR1* KO MG, which could contribute to phagocytic deficit and immune dysfunction. In order to test this hypothesis, we treated mFXS MG and *FMR1* KO MG with 25μM RAC1 inhibitor (iRAC) (NSC23766-[N6-[2-[[4-(diethylamino)-1-methylbutyl]amino]-6-methyl-4-pyrimidinyl]-2-methyl-4,6quinolinediaminetrihydro-chloride]) (Fig. 3C). Treatment with iRAC ameliorated the phagocytic deficit in mFXS-1 MG and *FMR1* KO MG (Fig. 3D, Supplementary movie 4). To test if iRAC treatment also ameliorated the enhanced actin polymerisation in microglial cells lacking FMRP, we measured the level of F actin during zymosan bioparticle phagocytosis at 60 minutes and 120 minutes in mFXS MG and *FMR1* KO MG. F actin staining and its quantification demonstrated that iRAC treatment significantly reduced the levels of F actin in both mFXS MG and *FMR1* KO MG at 60 min and 120 min during phagocytosis when compared to untreated cells (Figs. 3E-I).

Next, we investigated the status of the NF-κB pathway following iRAC treatment in mFXS MG and *FMR1* KO MG. iRAC-treated mFXS MG and *FMR1* KO MG demonstrated a significant reduction in the nuclear localisation of NF-κB when compared to untreated cells (Figs. 4J-M), suggesting suppression of NF-κB signaling. To test the functional consequence of the suppression of NF-κB signaling by iRAC treatment, we performed qPCR for proinflammatory genes, such as *IL1b* and *TNFa*, in iRAC-treated mFXS MG and *FMR1* KO MG. The qPCR results revealed significantly reduced expression of proinflammatory transcripts in iRAC-treated mFXS MG and *FMR1* KO MG when compared to untreated cells (Figs. 3N, 3O). Collectively, these findings indicate that inhibition of RAC1 signaling in microglial cells lacking FMRP ameliorates their phagocytic deficit and attenuates the production of proinflammatory cytokines.

**Figure 4:**
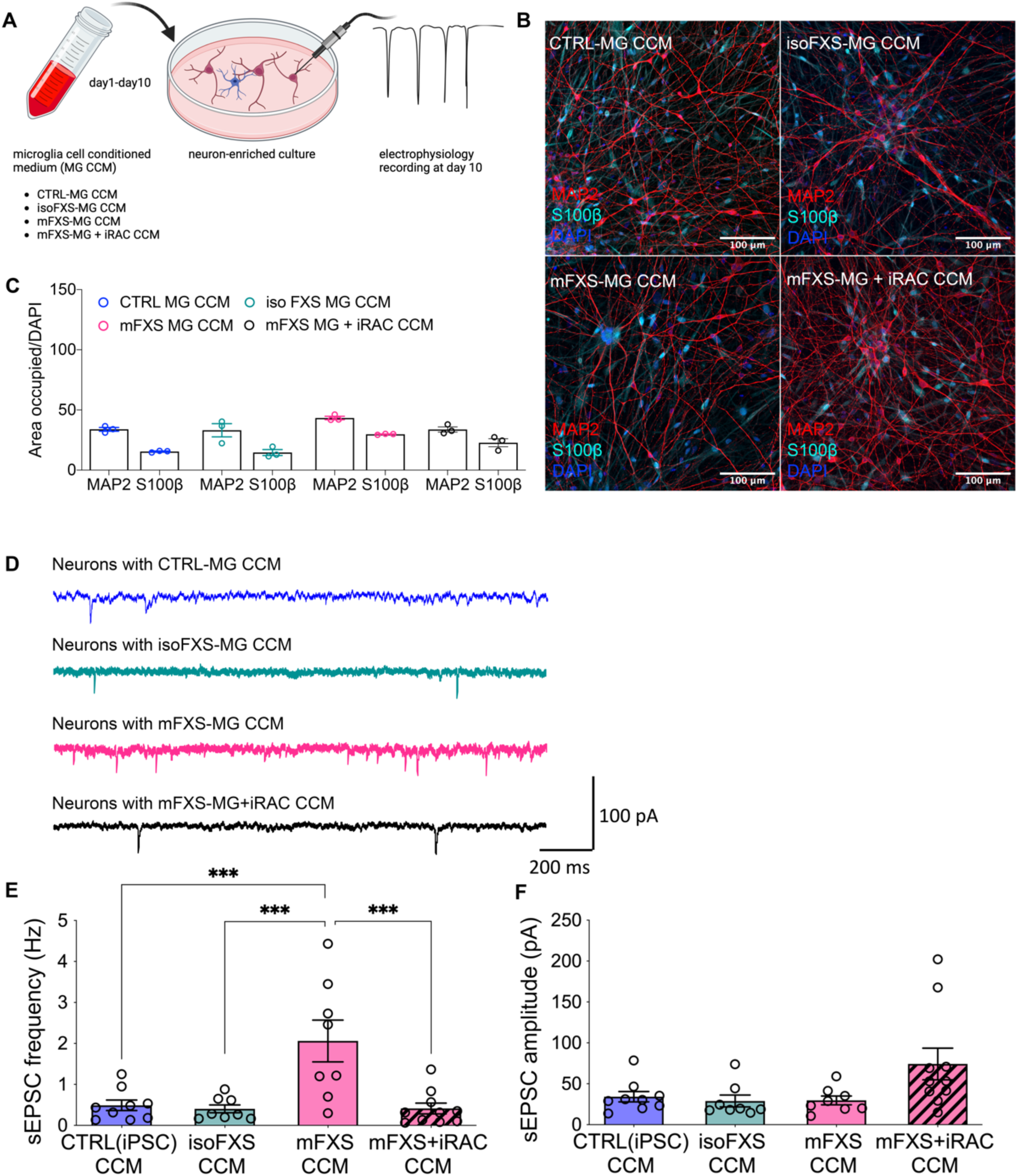
Cell conditioned medium derived from mFXS-MG prompts hyper-excitability in control human iPSC-derived neuronal culture. **(A)** Schematic representing the protocol for treating hiPSC neuronal cultures with cell conditioned medium (CCM) from CTRL-MG, isoFXS1-MG, mFXS1-MG and mFXS-MG1+iRAC for 10 days. Post 10 days, whole-cell patch clamp recordings were performed on individual neurons. **(B)** Representative immunofluorescent images showing staining for neuronal marker (MAP2) astrocyte marker (S100β) and nuclear marker (DAPI) in iPSC neuronal cultures treated with cell conditioned medium (CCM) derived from CTRL-MG, isoFXS1-MG, mFXS1-MG and mFXS1-MG+*i*RAC after 10 days. **(C)** Graph representing the quantification of MAP2 and S100β positive area normalized to DAPI in iPSC neuronal cultures treated with cell conditioned medium (CCM) derived from CTRL-MG, isoFXS1-MG, mFXS1-MG and mFXS1-MG+*i*RAC after 10 days. Data are represented as mean +/- SEM N=3; N represents the number of times experiments were performed using cells generated from independent neuronal differentiations. **(D)** Representative voltage-clamp recordings (V_H_ = −60 mV) of spontaneous excitatory postsynaptic currents (sEPSCs) from hiPSC derived cortical neurons treated with CCM from CTRL-MG (*blue trace*), isoFXS1-MG (*turquoise trace*), mFXS1-MG (*pink trace*) and mFXS1-MG+iRAC (*black trace*). **(E,F)** Comparison of mean sEPSC frequency **(E)** and mean sEPSC amplitude **(F)** of neurons treated with CCM from CTRL-MG, isoFXS1-MG, mFXS1-MG and mFXS1-MG+*i*RAC. Data represent mean ± SEM, N=3 statistical analysis was performed using on-way ANOVA with post-hoc Bonferroni’s test (*** p < 0.001). Each dot represent data from single cell. N represents number of times experiments were performed using cells generated from independent iPSC differentiations.

### Conditioned medium from mFXS-MG causes hyperexcitability in control iPSC derived neurons

It is well established that pro-inflammatory cytokines such as TNF-a and IL-1b can modulate neuronal excitability (27, 28). Given that neuronal hyperexcitability is a key disease feature in FXS (6) and loss of FMRP confers a pro-inflammatory state to microglial cells (Fig 2A-2D), we further investigated the impact of cell-conditioned medium (CCM) from mFXS MG on control iPSC-derived cortical neurons (Fig. 4A). To this end, we differentiated cortical neurons from control iPSCs employing an established protocol (29) and treated them with CCM for 10 days. We observed comparable proportions of neurons and astrocytes in the neuronal cultures treated with CCM derived from CTRL-MG, isoFXS-MG, mFXS-MG, and mFXS-MG+iRAC (Figs 4B, C). Whole cell patch clamp based electrophysiological recording revealed that visually identified neurons across all experimental groups were able to fire trains of action potentials following current injection; intrinsic properties for neurons are summarised in Supplementary Table 2. To assess changes at the level of neuronal network activity we measured their spontaneous excitatory post-synaptic currents (sEPSCs) and sEPSCs are represented by heterogeneous inward currents of variable amplitudes. We noted significantly increased sPSCs in neuronal cultures treated with CCM from mFXS MG when compared to CCM from isoFXS MG and CTRL MG (Figs. 4D-4F), suggestive of an increased network excitability. In the presence of CCM from mFXS MG, the neurons exhibited higher input resistance (Supplementary Table 2), which could contribute to the increased network excitability. Interestingly, there was no significant change in the amplitudes of neuronal sEPSCs across the different CCM-treated conditions (Fig. 4E). Importantly, CCM from iRAC-treated mFXS MG, ameliorated the enhanced sEPSC frequency seen in neuronal cultures treated with CCM from mFXS MG CCM treated neuronal cultures (Figs. 4D-F). Collectively this experiment highlights that CCM from mFXS MG prompts network hyperexcitability in human cortical neurons, an effect that could be prevented by iRAC treatment of mFXS MG.

## Discussion

Here we provide evidence for both cell autonomous and non-cell autonomous role of microglial cells in the pathophysiology of FXS. Using a range of human cellular platforms, we found that loss-of-function of *FMR1* leads to a proinflammatory immune state and phagocytic deficits in microglia through increased RAC1 signaling.

To the best of our knowledge, this is the first report of phagocytic deficit and immune dysfunction in human microglia carrying *FMR1* mutation. Notably, other autism causing mutations such as loss-of-function of either X-linked *Mecp2 or Nlgn4* leads to phagocytic deficit in microglia (30, 31). It is well established that FXS patients and *fmr1*-/- mice display increased number of dendritic spines and synapses (32–34), which could result from microglial dysfunction as microglia have been suggested to play a pivotal role in remodelling dendritic spines during development (35). Chimeric rodent models or 3D organoid models could be further used to investigate role of diseased microglia during dendritic spine development in FXS (36, 37).

Our finding highlighting the proinflammatory profile of FXS microglia are in line with previous observations indicating increased levels of proinflammatory cytokines in the blood of FXS (14) and patients with autism (38–40). To this end, Takada et al have shown that increased level of proinflammatory cytokines produced from differentiated macrophages from the peripheral blood of patients with autism can negatively impact dendritic outgrowth of iPSC derived neurons (41). Notably, the impact of microglial activation on neuronal morphology, function and stress associated behaviour has been described in many neurodevelopmental disorders including Rett’s syndrome, schizophrenia and Downs syndrome (42–44) and it is of interest that some clinical trial data suggest that suppression of inflammation may improve cognition in schizophrenia patients and autism (45–47).

Our finding that mFXS-MG contribute to network hyperexcitability of cortical neurons, is consistent with existing literature demonstrating that proinflammatory cytokines can impact both spontaneous excitatory post synaptic currents (sEPSC) and mini excitatory postsynaptic currents (mEPSC) (28), which could be mediated through direct interaction of cytokines with receptor and ion channels in neurons (48). Interestingly, it is well known that FXS neurons are hyperexcitable and although a range of causal mechanisms such as synaptic, and circuit-level dysfunctions have been proposed to explain hyperexcitability (6), our data suggest an additional mechanism by which proinflammatory microglial cells could contribute to neuronal hyperexcitability in FXS. However, to better understand the molecular mechanisms underlying neuronal hyperexcitability, transcriptomics or proteomics could be performed with neuronal cultures treated with diseased microglial conditioned medium.

We observe that mFXS-MG display activated RAC1 signaling, which corroborates with the existing literature on FXS neurons demonstrating that loss of function of FMRP leads to an activated RAC1-COFILIN axis, and aberrant actin polymerisation (49, 50). Given that, RAC1 signaling is also implicated in phagocytosis and immune regulation (51, 52), we suggest that activated RAC1 signaling in mFXS-MG underlies (1) the phagocytic deficit through aberrant actin dynamics and (2) the proinflammatory state of microglial cells through the activated NF-κB pathway. Our findings related to increased level of phospho COFILIN in mFXS-MG are in line with experimental models of COFILIN inactivation in mammalian cells, which also display phagocytic deficits through altered actin dynamics (53). Furthermore, our findings highlighting reduced activation of NFκB in iRAC treated mFXS-MG (Figs 3L-3O) are consistent with previous reports demonstrating that, inhibition of RAC1 pathways either through *Rac1* siRNAs or NSC23766, significantly reduces NF-κB activity (52).

While treatment with RAC1 inhibitor NSC23766, has been shown to alleviate cognitive impairments in *Fmr1*^-/-^ mice through improvement of neuronal spine cytoarchitecture and synaptic deficits (54), our current findings highlight an immunomodulatory role of RAC1 inhibitors targeting human microglial cells in FXS. Taken together, our data suggest that loss-of-function of *FMR1* leads to microglial activation and phagocytic deficits via activated RAC1 signaling. Exposure of human control iPSC-derived cortical neurons to conditioned medium from activated mFXS MG results in neuronal hyperexcitability. Importantly, pharmacological suppression of microglial RAC1 signaling ameliorates neuronal hyperexcitability, collectively suggesting that RAC1 signaling could emerge as a potential therapeutic target in FXS.

## Materials and methods

### Generation of FXS isogenic iPSC line

A control iPSC line (CTRL), mFXS-1 patient iPSC lines harbouring the CGG repeat expansion, its respective isogenic control lines (isoFXS-1), *FMR1* null line and a control embryonic stem cell control line used in this study were generated with full Ethical/Institutional Review Board approval at the University of Edinburgh, as previously reported by our group (55). The mFXS-2 iPSC line carrying the CGG repeat expansion mutation was gene corrected using CRISPR/Cas9 genome editing as described in previous publication by our group (55) (Supplementary Fig 1). Briefly, iPSCs were dissociated into single-cell suspension with 1X Accutase (Sigma-Aldrich). 8 × 10^5^ cells were transfected with 2 mg Cas9-gRNA-1 plasmid, 2 mg Cas9-gRNA-2 plasmid and 1 mg eGFP-puromycin resistance plasmid using the Amaxa 4D-Nucleofector system (Lonza) according to the manufacturer’s instructions with a pulse load of CA137. Cells were plated down onto MatrigelTM (Corning) coated plates in E8 medium with 10 mM ROCK inhibitor (Tocris). After 24 h, transfected cells were selected for with addition of 1 mg/mL puromycin (Sigma-Aldrich) for a further 24 h. Clonal analyses were performed on manually picked individual clones and screened for successful gene-editing via immunofluorescence for FMRP and PCR flanking repeat expansion mutation. Successful gene editing clone was determined by re-expression of FMRP and presence of positive PCR amplicon. Successful gene-edited clones were further validated using repeat primed PCR, using AmplideX FMRI PCR kit from Asuragen, which showed negative for gene-edited clone.

### Differentiation and characterisation of microglia (hiPSC-MG) from human iPSC

A control iPSC line (CTRL), two FXS patient iPSC lines (mFXS-1, mFXS-2) harbouring the CGG repeat expansion and their respective isogenic control lines (isoFXS-1, isoFXS-2) alongside control human embryonic stem cell (CTRL-ES) and *FMR1* knock out line (FMR1KO) used in this study were maintained in Essential 8 Medium (E8, Gibco) in plates coated with Matrigel Growth Factor Reduced Basement Membrane Matrix and were passaged at 80% confluence with 1 mg/ml collagenase and 0.5 mg/ml dispase (Thermo Fisher Scientific). All iPSCs were confirmed to be sterile, mycoplasma-free and G-banding was performed at regular intervals to exclude the presence of any acquired chromosomal abnormalities in the iPSCs. Details of the cell lines used are summarised in Supplementary Table 1.

Human microglia-like cells (hiPSC-MG) were generated following a protocol previously described by our group (25). Briefly, myeloid progenitors expressing CX3CR1, CD11b and CD45 were derived from mesodermally-patterned embryoid bodies cultured in myeloid precursor media containing 100 ng/ml M-CSF (Peprotech) and 25 ng/ml IL-3 (Gibco) for 4-6 weeks. These myeloid precursors were then differentiated to microglia-like cells (hiPSC-MG) on poly-D-lysine treated gelatinised plates (0.1%) in the presence of 100 ng/ml IL-34 (Peprotech), 10 ng/ml GM-CSF (Peprotech) and line-matched neural precursor cell conditioned medium (NPC-CM), which was supplemented in an increasing gradient (10%-50%) every day from Day 7. Details of reagents used are summarised in Supplementary Table 3.

### Immunocytochemistry

hiPSC-MG grown on coverslips were fixed in 4% paraformaldehyde (PFA; Agar scientific) in phosphate-buffered saline (1x PBS) for 10 min at room temperature. After fixation, cells were washed three times with 1x PBS, permeabilised with 0.3% Triton-X100 (Thermo Fisher Scientific) in 1x PBS for 5 min and blocked with 6% BSA (Europa Bioproducts) in 1x PBS for 1hr at room temperature. Incubation with primary antibodies for two hours was followed by addition of appropriate secondary antibodies. The cells were then washed three times with 1xPBS, counterstained with 4′,6-diamidino-2-phenylindole (DAPI; Thermo Fisher Scientific) and mounted on to glass slides in FluorSave™(Millipore). The slides were observed and imaged using a Zeiss LSM 710 confocal microscope. Details of antibodies used in the study are summarised in a resource table (Supplementary Table 3)

### Quantification of immunofluorescence images using Cell Profiler software

The image analysis was performed using the cell profiler software (version 4.2.1). Z-stacked images were acquired using confocal microscopy (Zeiss LSM 710) and saved as .tiff files after separation into different channels using ImageJ Fiji software (version 2.14.0). Subsequently, the images were imported into cell profiler for analysis. The analysis involves classifying objects into primary, secondary, and tertiary. To quantify the area occupied or the intensity of the immunofluorescence staining of the desired object, a custom mask is designed using the in-built modules in the Cell profiler. The objects were segmented based on a Global threshold strategy to separate the two classes of pixels (foreground and background). The threshold correction factor was set between 1 and >1 making the threshold more stringent, and the threshold smoothing factor was set at 1.3488 to improve the uniformity, removing holes, and noise in the image. The method to distinguish the clumped objects was based on intensity and the objects that reside outside the diameter range were discarded.

### RNA extraction, reverse transcription and polymerase chain reaction (PCR)

Cells were harvested and RNA was extracted using RNeasy Mini Kit (Qiagen). Contaminating DNA was removed using RNase-Free DNase Set (Qiagen). The RNA was reverse transcribed using RevertAid RT Reverse Transcription Kit (Thermo Fisher Scientific) according to the manufacturer’s protocol. Quantitative real-time PCR (qRT-PCR) was performed with DyNAmo ColorFlash SYBR Green Master Mix (Thermo Fisher Scientific) on a CFX96™ Real Time PCR System (BioRad). Primer sequences are reported in earlier publication (25)

### Flow cytometry

The myeloid precursors were collected from the supernatant, centrifuged, washed with 1x PBS and were treated with 1% Fc block (Miltenyi Biotec) in 1x PBS for 10 min at room temperature, before incubating them with primary antibodies for one hour on ice. Samples were then washed twice with 1x PBS and transferred onto round-bottom polystyrene tubes (BD Falcon) until assessment. The stained cell samples were analysed using a FACS LSR Fortessa (4 lasers) flow cytometer (BD Biosciences), and the data were processed using FlowJo software.

### High-throughput imaging and analysis platform for the quantification of microglial phagocytosis

Microglial differentiation was performed in a 96-well black/clear bottom plate with optical polymer base (Thermo Scientific™) for 12 days. pHrodo-conjugated zymosan bioparticles (Thermo Fisher Scientific) were added to the hiPSC-MGs at a concentration of 0.5 mg/ml and the plate was immediately loaded into ImageXpress Micro 4 High-Content Imaging System, a high-throughput live imaging system maintained at 37 degrees C and 5% CO_2_ to monitor a real-time uptake of zymosan bioparticles by the hiPSC-MGs. Prior to loading, hiPSC-MG were co-stained with Hoechst 33342 (Thermo Fisher Scientific). The cells were imaged using a 20X objective at an interval of 15 min for 2.15 h (135 min). All lines reported in this study had a minimum of 3 biological replicates.

In order to determine the phagocytic index of individual genotypes (mutant, isogenic and control) images were analysed using MetaXpress software. Nuclei were first detected using a fluorescence threshold to separate background pixels from Hoechst-stained objects. Objects were then classified as nuclei based on their size. The nuclei were then expanded by 50 pixels and within this area we used a contrast threshold to identify the borders of pHrodo objects. From each image we calculated the number of nuclei (Hoechst-stained objects), the total intensity of pHrodo objects, and the data was plotted as intensity/microglial cell (MG) after normalising the total intensity of pHrodo objects with total number of nuclei.

### Forward phase cytokine microarray

Conditioned medium per genotype was harvested from 2 × 10^6^ microglial cells per dish. The cell conditioned media was spun at 2000 RPM for 10 min to remove cellular debris. 64 capture antibodies were printed as 64 sub-arrays with 4 replicates per target on a single nitrocellulose-coated glass slide (Supernova Grace Biolabs) using a Quanterix-2470-microarrayer and 185uM printing pins as previously described (56). Cell conditioned medium samples were each incubated with identical 64 sub-array slides in order to generate four readout values per sample. After blocking, sample incubation and repeated wash steps, arrays were incubated with a specific detection antibody, which was biotin labelled. A final incubation with fluorescently labelled streptavidin generated the signal for all samples and this was quantified using an InnoScan710IR scanner (Innopsys) and analysed using Mapix software (Innopsys). After collection of conditioned medium, the cells attached on the plates were washed once with 1x PBS and lysed in RIPA lysis buffer for performing a protein estimation which was used for normalization of the cytokine array data.

### Western blot analysis

Western blot analysis was carried out according to an established protocol (57). Briefly, the cells were lysed in lysis buffer (20 mM Tris-Cl pH 7.4, 1 mM EDTA, 150 mM NaCl, 1% Triton X-100, 1 mM EGTA, 1mM phenylmethylsulfonyl fluoride). The protein concentration was calculated using the BCA Protein Assay Reagent (23225, Pierce™ Thermo Fisher Scientific). Protein samples were separated by SDS-PAGE (Thermo Fisher Scientific) and transferred to 0.25 micron polyvinylidene difluoride (PVDF) membranes (Millipore, Bedford, MA, USA). The membranes were blocked with 5% skimmed milk and incubated at 4 °C overnight with primary antibodies followed by addition secondary antibodies with IRDye 680RD and IRDye 800CW (Li-COR Bioscience, USA) respectively for 1 hr at room temperature. Following washing, the blot was imaged using Li-COR Odyssey FC infrared system.

### Assessment of F actin and G actin

F actin and G actin fractionation was performed by adopting an established protocol(58). Briefly, microglial cells across genotypes were washed three times in 37 °C PBS before lysis with actin stabilization buffer (0.1 M PIPES pH 6.9, 30% glycerol (vol/vol), 5% DMSO (vol/vol),1 mM MgSO4, 1 mM EGTA, 1% Triton X-100 (vol/vol), 1 mM ATP, complete protease inhibitor and phosphatase inhibitor. The lysis was performed at 37 °C for 10 min. Cells were dislodged by scraping and the entire extract was centrifuged at 4 °C for 120 min in an ultracentrifuge (Beckmann, rotor TLA-120.1) at 100,000g. The supernatant containing G-actin was recovered and the pellet containing F-actin collected separately and resuspended with RIPA buffer. The total protein concentration was quantified by BCA (Pierce BCA Protein Assay Kit) and equal protein concentrations were used for western blotting.

### RAC activation assay

PAK-PBD protein specifically recognizes and binds the active, GTP-bound, forms of the RAC. We examined RAC activity using the PAK-PBD protein beads coupled to a coloured glutathione Sepharose matrix. 1mg of cell lysates across all genotypes were incubated with PAK-PBD beads for 1hr at 4°C on a rocker. Following incubation, beads were pelleted by centrifugation (14,000g for 15 s at 4°C), and the supernatant was discarded. The pelleted beads were washed with lysis buffer three times, resuspended in 100μl of 4× Laemmli buffer, boiled for 5 min, and were used for Western blot analysis with RAC1 antibody.

### Generation of cortical neurons from hiPSC and treatment with microglia conditioned medium

Human iPSC-derived cortical neuronal precursor cells (NPC) were generated as described previously (29). NPCs were plated in Default media comprising Advanced-DMEM/F12, 1% P/S, 0.5% Glutamax, 0.5% N2, 0.2% B27, 2 µg/mL Heparin (Sigma) on poly-D-lysine (Sigma), laminin (Sigma), fibronectin (Sigma) and matrigel coated coverslips for differentiation and fed twice a week. Default media was supplemented with 10 µM forskolin (Tocris) in weeks 2 and 3. From week 4 onwards forskolin was removed and Default media was supplemented with 5 ng/mL BDNF and 5 ng/mL GDNF and the cells were cultured till week 6.

From week 7, 20% microglial conditioned medium derived from CTRL-MG, isoFXS-MG, mFXS-MG and mFXS-MG+iRAC were added onto the neuronal cultures every other day for 10 days. The conditioned medium was centrifuged at 2000 RPM for 10 min to remove any cellular debris before adding onto the neuronal cultures. The neurons were patched at week 8 for electrophysiology recording.

### Electrophysiology

Whole-cell patch clamp recordings were performed using similar protocols as described previously (55). Briefly, coverslips containing cultures of hiPSC-derived cortical neurons were transferred to a recording chamber, which was continuously perfused with an external recording solution that had the following composition (in mM): NaCl 152, KCl 2.8,HEPES 10, CaCl_2_ 2, glucose 10, pH 7.3–7.4 (300–320 mOsm). Patch-pipettes, fabricated from thick-walled borosilicate glass, with a resistance of 3-4 MΩ were filled with an internal solution comprising (in mM): K-gluconate 155, MgCl_2_ 2, HEPES 10, Na-PiCreatine 10, Mg_2_-ATP 2, and Na_3_-GTP 0.3, pH 7.3 (280–290 mOsm). Spontaneous excitatory postsynaptic currents were recorded at a voltage of −60 mV. All the recordings were performed using a MultiClamp 700B amplifier (Molecular Devices, San Jose, CA). Data were sampled at 2.8 kHz and digitized at 10 kHz via Digidata 1550A. Subsequent offline analysis was performed using Clampfit 10.7 software.

## CRediT Contribution

Poulomi Banerjee (PB): Conceptualization, Validation, Investigation, Visualization, Methodology, writing — original draft, Project administration, Writing – review and editing Shreya Das Sharma (SDS): Methodology, Investigation, Formal Analysis, Writing – original draft

Karen Burr (KB): Methodology, Investigation,

Kimerley Morris (KM) Investigation

Tuula Ritakari (TR)-Methodology, Investigation

Paul Baxter (PB): Investigation

James D Cooper (JDC): Investigation

Alessandra Cardinalli (AC): Formal Analysis

Srividya Subash (SS): Investigation, Formal Analysis

Evdokia Paza (EP): Investigation

David Story (DS): Resources (administrative support)

Sumantra Chattarji (SChattarji): Project administration, Writing - Review & Editing

Peter C Kind (PCK): Project administration, Writing - Review & Editing

Neil O Carragher (NC): Resources (cytokine assay development platform), Writing – review and editing

Bhuvaneish T Selvaraj (BTS): Conceptualization of gene editing strategy, Methodology, Project administration, Writing – review and editing

Josef Priller (JP): Conceptualization, Supervision, Project administration, Writing - Review & Editing

Siddharthan Chandran (SChandran): Conceptualization, Resources, Supervision, Project administration, Visualization, Writing - Original Draft, Writing - Review & Editing, Funding acquisition

## Acknowledgements

Flow cytometry data were generated within the Flow Cytometry and Cell Sorting Facility in MRC Centre for Regenerative Medicine. Cytokine microarray data was generated by Alison Munro and Camilla Drake within the HTPU Microarray services facility in the Institute of Genetics and Cancer. Authors acknowledge the many stimulating discussions with scientists based in National Centre for Biological Sciences, Tata Institute of Fundamental Research, and Centre for Brain Development and Repair, Institute for Stem Cell Biology and Regenerative Medicine, Bangalore India through a collaborative research programme which helped frame this study. Some of the illustrations in the manuscript were created with BioRender.com.

For the purpose of open access, the authors have applied a CC-BY public copyright licence to any Author Accepted Manuscript version arising from this submission

---

**Supplementary Fig 1:**
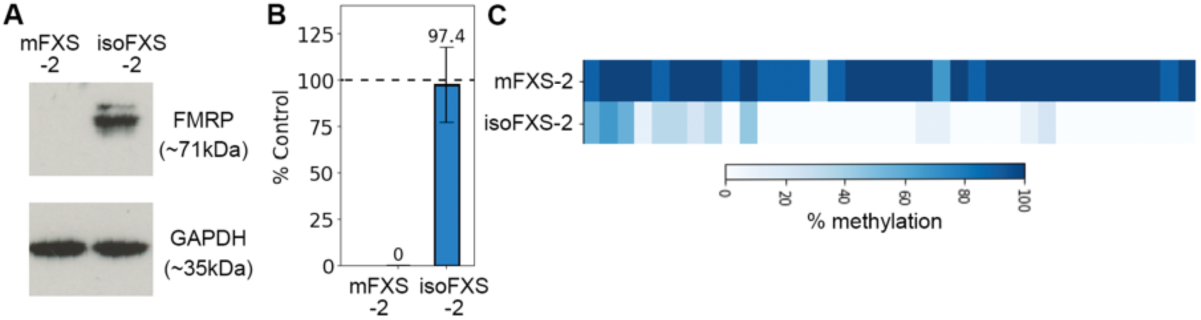
Validation of isoFXS-2 induced pluripotent stem cells. **(A,B)** Western blot and its quantification depicting loss of FMRP in mFXS-2 iPSC line and re-expression of FMRP in isoFXS-2 iPSC line post gene editing. **(C)** Bisulfite sequencing of the *FMR1* promoter showing its demethylation in isoFXS-2 iPSC line

**Supplementary Fig 2:**
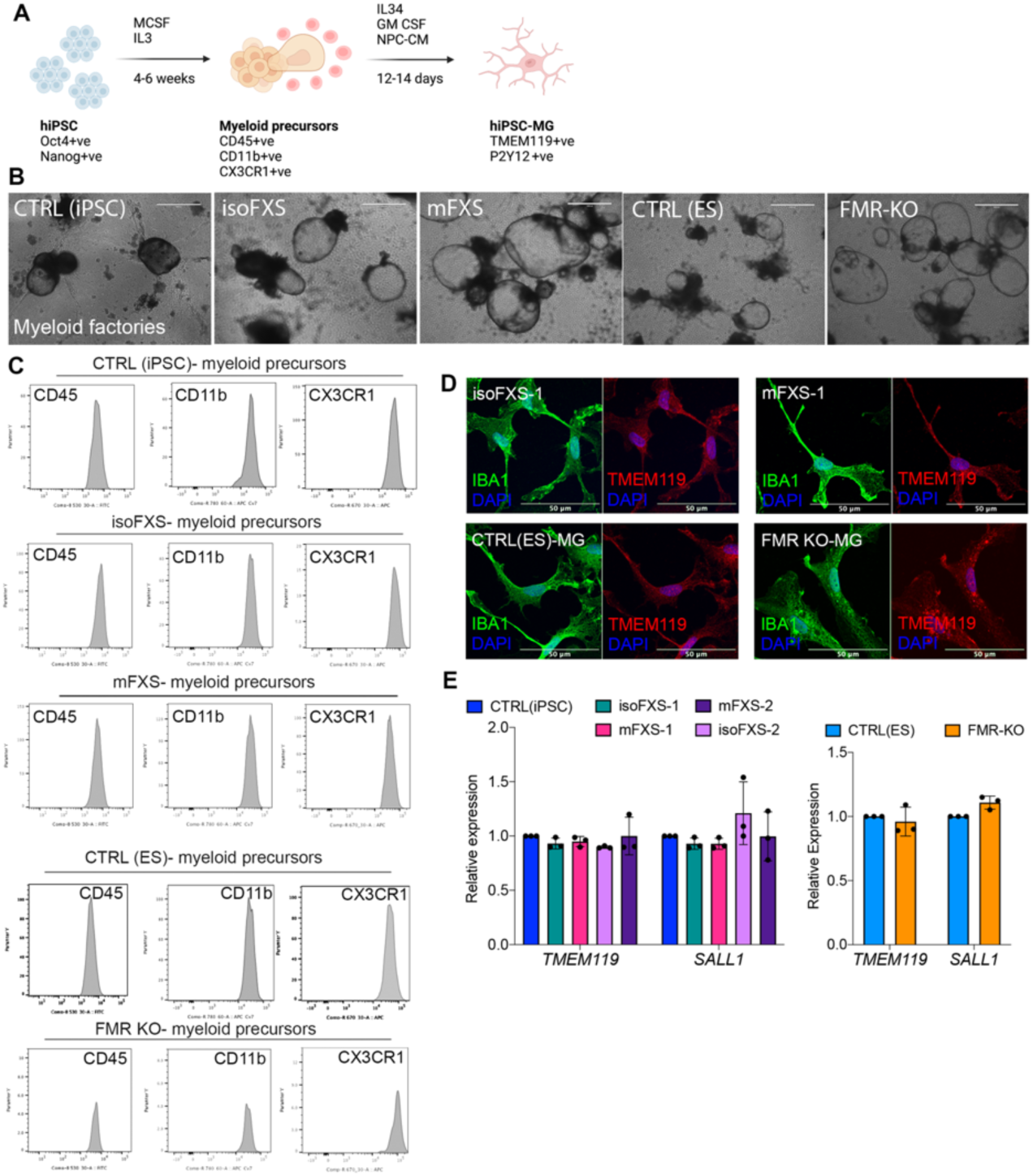
Generation and characterisation of mFXS-MG, isoFXS-MG, CTRL(iPSC), *FMR1* KO-MG and CTRL (ES)-MG. **(A)** Schematic representing experimental paradigm to differentiate microglial-like cells from induced pluripotent stem cells **(B)** Phase contrast images showing the formation of myeloid factories from CTRL(iPSC), isoFXS, mFXS, and CTRL(ES), *FMR1* KO, Scale bar=200μm **(C)** Myeloid precursors produced by the CTRL (iPSC), isoFXS1, mFXS1, CTRL(ES), *FMR1* KO myeloid factories show comparable profile for key myeloid precursor markers-CD45, CX3CR1 and CD11b **(D)** Representative immunofluorescence images of isoFXS1-MG, mFXS1-MG, CTRL (ES)-MG and *FMR1* KO-MG demonstrating comparable staining of microglial markers: TMEM119 (red) IBA-1 (green) **(E)** Bar graphs representing comparable expression of microglia signature genes (*TMEM119, SALL1*) in CTRL(iPSC)-MG, isoFXS1-MG, mFXS1-MG, isoFXS2-MG, mFXS2-MG CTRL(ES)-MG and *FMR1* KO-MG.

**Supplementary Fig 3:**
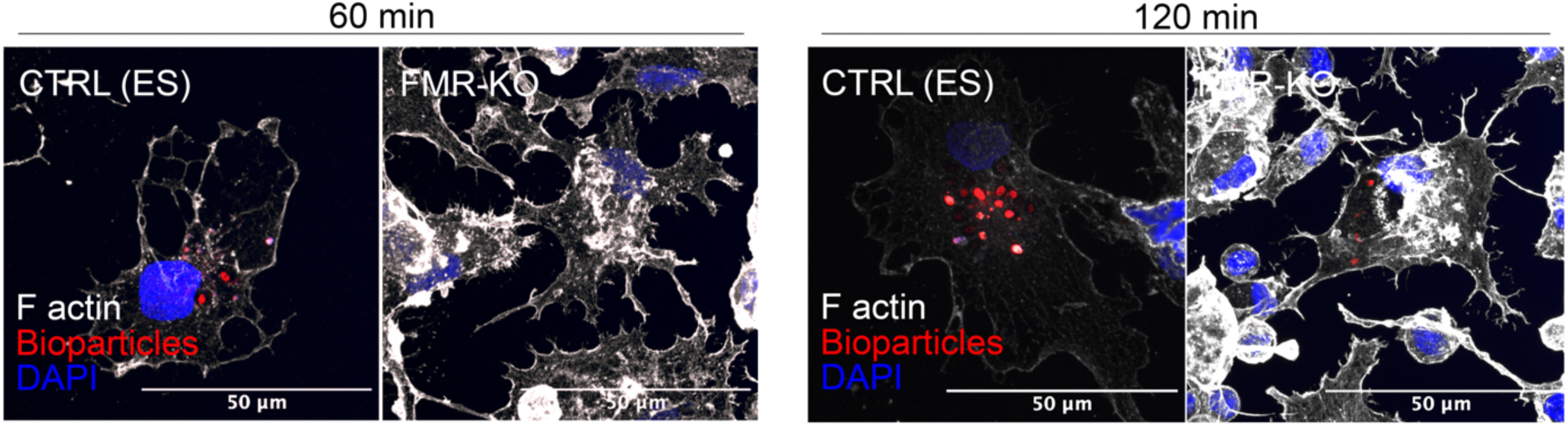
Localisation of F actin in *FMR1* KO-MG and CTRL (ES)-MG during phagocytosis of zymosan bioparticle. Representative images of immunofluorescence staining with phalloidin showing aberrant F actin in *FMR1* KO-MG when compared to CTRL (ES)-MG at 60-minutes (left panel) and 120-minutes (right panel) during phagocytosis of zymosan bioparticles (red).

**Supplementary Table 1:**
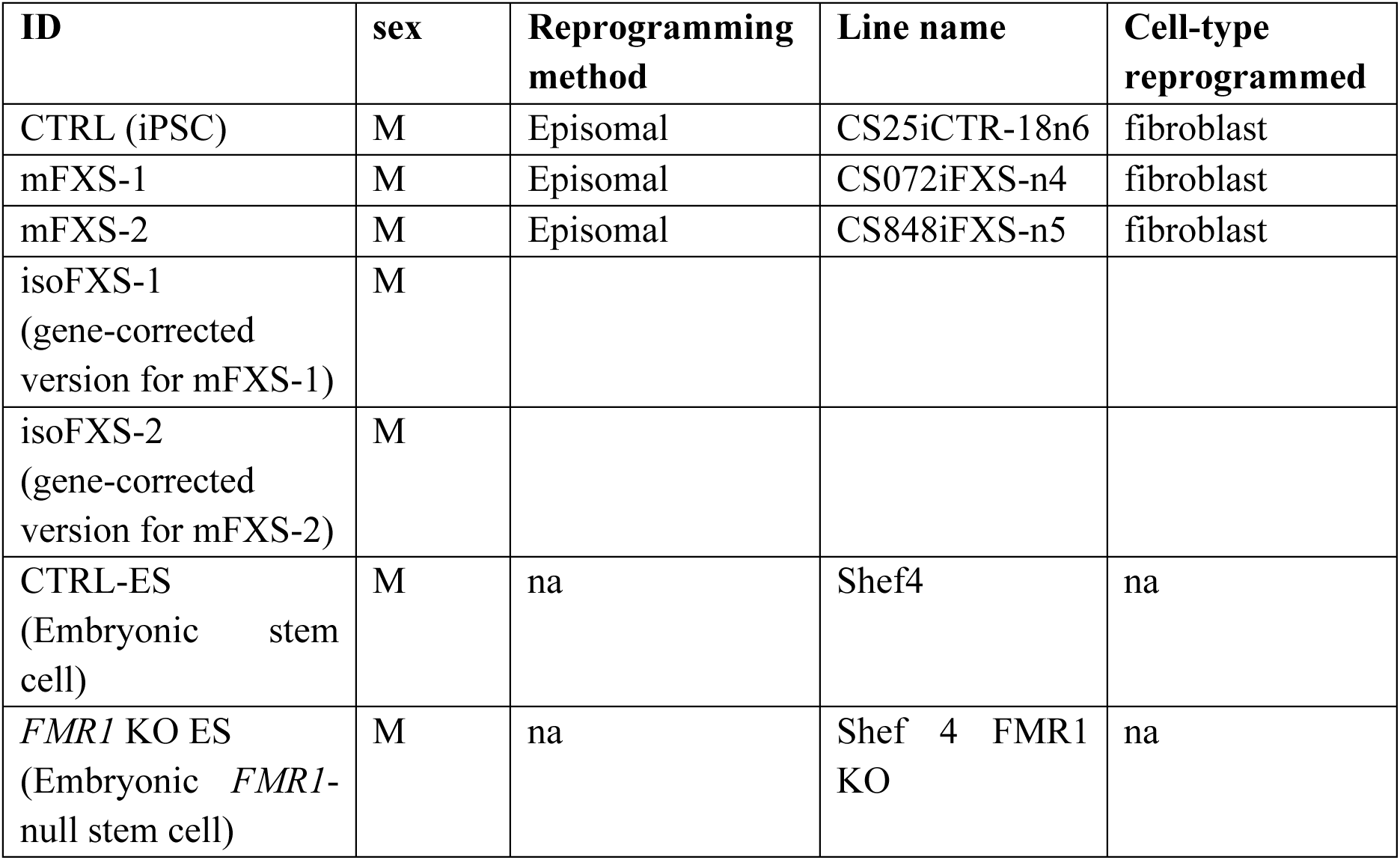
Details for pluripotent lines used in the study.

**Supplementary Table 2:** Intrinsic property of iPSC-neurons following treatment with cell conditioned medium (CCM) generated from CTRL-MG, isoFXS-1MG, mFXS1-MG and mFXS-MG+iRAC.

|  | CTRL-MG CCM | isoFXS1-MG CCM | mFXS1-MG CCM | mFXS1-MG+iRAC CCM |
| --- | --- | --- | --- | --- |
| Number of cells | 9 | 8 | 8 | 10 |
| | Mean ± SEM | Mean ± SEM | Mean ± SEM | Mean ± SEM |
| Resting membrane potential (mV) | -62.98 ± 2.2 | -57.6 ± 2.35 | -54.92 ± 3.08 | -58.1 ± 2.29 |
| Input Resistance (GΩ) | 1.18 ± 0.15 | 1.15 ± 0.11 | 1.79 ± 0.19 | 1.12 ± 0.07 |
| Capacitance (pF) | 38.71 ± 6.07 | 30.85 ± 2.25 | 31.44 ± 4.42 | 34.84 ± 5.73 |
| Rheobase (pA) | 13.89 ± 1.82 | 20.56 ± 3.16 | 7.5 ± 1.34 | 14.5 ± 2.03 |
| Spike amplitude (mV) | 57.2 ± 3.67 | 49.78 ± 3.08 | 52.52 ± 2.57 | 59.63 ± 4.48 |
| Spike half-width (ms) | 5.21 ± 0.92 | 6.07 ± 0.63 | 5.96 ± 1.26 | 5.25 ± 1.03 |
| Max. no. of Action Potential (AP) fired | 2.8 ± 0.52 | 1.7 ± 0.3 | 2.4 ± 0.52 | 2.9 ± 0.45 |

**Supplementary Table 3:** (Resource Table)

| **Name** | **cat codes.** | **Company** | **working concentration** |
| --- | --- | --- | --- |
| Essential 8™ Medium | A1517001 | Gibco | NA |
| Collagenase Type IV | 17104019 | Thermo Fisher Scientific | 1mg/ml |
| DispaseII | 17105041 | Thermo Fisher Scientific | 0.5mg/ml |
| Rock Inhibitor | 1254 | Tocris | 10μM |
| X-VIVO 15 Serum-free Hematopoietic Cell Medium | BE02-060F | Lonza | NA |
| Advanced DMEM/F12 | 12634028 | Thermo Fisher Scientific | 0.5X |
| Neurobasal | 21103049 | Thermo Fisher Scientific | 0.5X |
| GlutaMAX™ Supplement | 35050061 | Thermo Fisher Scientific | 1X |
| MEM Non-Essential Amino Acids Solution (100X) | 11140050 | Thermo Fisher Scientific | 1X |
| N-2 Supplement (100X) | 17502048 | Thermo Fisher Scientific | 1X |
| β-Mercaptoethanol | 31350-010 | Thermofisher Scientific | 100μM |
| Antibiotic Antimycotic Solution (100×) | A5955 | Merck | 1X |
| Human MCSF | 300-25 | Peprtech | 100ng/ml |
| Human IL 3 | PHC0035 | Gibco | 25ng/ml |
| Human GMCSF | 300-03 | Peprtech | 10ng/ml |
| Human IL34 | 200-34 | Peprtech | 100ng/ml |
| Human BMP4 | 120-05ET | Peprtech | 50ng/ml |
| Human VEGF | 100-20 | Peprtech | 50ng/ml |
| Human SCF | 300-07 | Peprtech | 25ng/ml |
| Poly-D-lysine hydrobromide | P6407 | Sigma | 0.1mg/ml |
| Gelatin | G1393 | Merck | 0.1% |
| DPBS | 14040133 | Invitrogen | NA |
| Accutase | A6964 | Merck | 1X |
| B27 with Vit A | 17504044 | Thermo Fisher Scientific | 1X |
| BDNF | 248-BDB | R & D Systems | 10 ng/ml |
| GDNF | 212-GD/CF | R & D Systems | 10 ng/ml |
| FGF-2 | 450-33 | PeprTech | 10ng/ml |
| Laminin | L2020 | Sigma-Aldrich | 5.0 µg/ml |
| Fibronectin | F2006 | Sigma-Aldrich | 20µg/ml |
| Matrigel Growth factor Reduced | 354230 | Corning | 1:30 for IPSC, 1:100 for c neurons |
| PAK-PBD Beads | PAK02-A | Cytoskeleton, Inc | 20 µg beads for 1mg protein |
| Alexa Fluor™ 555 Phalloidin | A34055 | Thermo Fisher Scientific | 1:1000 |
| NFκB antibody | ab16502 | Abcam | 1:300 |
| pHrodo™ Red Zymosan Bioparticles™ Conjugate for Phagocytosis | P35364 | Thermo Fisher Scientific | As per manufacturer's instructions |
| MAP2 antibody | A85363 | antibodies.com | 1:1000 |
| S100β antibody | ab52642 | Abcam | 1:500 |
| TMEM119 antibody | ab185333 | Abcam | 1:200 |
| IBA 1 antibody | ab5076 | Abcam | 1:500 |
| Phospho-Cofilin (Ser3) (77G2) | 3313 | Cell Signaling Technology | 1:2000 |
| Total anti-Cofilin | 5175 | Cell Signaling Technology | 1:1000 |
| Anti-FMRP antibody | ab17722 | Abcam | 1:1000 |
| RAC inhibitor<br>NSC23766- [N6-[2-<br>[[4-(diethylamino)-1-methylbutyl]amino]-<br>6-methyl-4-pyrimidinyl]-2-methyl-4,6quinolinediaminetr<br>ihydro-chloride] | 553502 | Sigma-Aldrich | 50µM |
| RDye® 800CW Goat<br>anti-Rabbit IgG (H +<br>L) | 926-32211 | LI-COR | 1:10,000 |
| IRDye® 680RD Goat<br>anti-Mouse IgG<br>Secondary Antibody | 926-68070 | LI-COR | 1:10,000 |
| FITC anti-CD45<br>(clone 2D1) | 304005 | Biolegend | 1:250 |
| APC anti-CD11b<br>(clone ICRF44) | 301309 | Biolegend | 1:300 |
| APC anti-human<br>CX3CR1 Antibody | 341609 | Biolegend | 1:250 |
| PVDF Transfer<br>Membrane, 0.2 µm | 88520 | Thermo Fisher Scientific |  |
| ROCHE cOmplete™<br>Protease Inhibitor<br>Cocktail | 12352200 | Millipore |  |
| 9cm² petri dishes | PDS-149-050F | Thermo Fisher Scientific |  |
| 500ml Rapid Flow unit | 10199-655 | Thermo Fisher Scientific |  |
| Anti-Actin antibody | A3853 | Sigma-Aldrich | 1:10,000 |
| **Software** |  |  |  |
| GraphPad Prism 7 | GraphPad Software | www.graphpad.com |  |
| Clampfit 11.1 | Molecular devices | www.moleculardevices.com |  |
| Cell Profiler | Cell Profiler software | https://cellprofiler.org |  |
| ImageXpress | Molecular devices | www.moleculardevices.com |  |

**Supplementary movie: 1**

Video showing phagocytosis of zymosan bioparticles (in red) in CTRL-MG over 2.15h

Bioparticles are pH sensitive, they will appear colourless in Time Point 0, however following internalisation, they will appear red inside the microglial cells.

**Supplementary movie :2**

Video showing phagocytosis of zymosan bioparticles (in red) in isoFXS-MG over 2.15h

Bioparticles are pH sensitive, therefore, they will appear colourless in Time Point 0, however following internalisation, they will appear red inside the microglial cells.

**Supplementary movie: 3**

Video showing phagocytosis of zymosan bioparticles (in red) in mFXS-MG over 2.15h

Bioparticles are pH sensitive, therefore, they will appear colourless in Time Point 0, however following internalisation, they will appear red inside the microglial cells.

**Supplementary movie: 4**

Video showing phagocytosis of zymosan bioparticles (in red) in iRAC treated mFXS-MG over 2.15h

Bioparticles are pH sensitive, therefore, they will appear colourless in Time Point 0, however following internalisation, they will appear red inside the microglial cells.

